# A magnetic pulse does not affect free-flight navigation behaviour of a medium-distance songbird migrant in spring

**DOI:** 10.1101/2022.04.28.489840

**Authors:** Thiemo Karwinkel, Michael Winklhofer, Lars Erik Janner, Vera Brust, Ommo Hüppop, Franz Bairlein, Heiko Schmaljohann

## Abstract

Current evidence suggests that migratory animals extract map information from the geomagnetic field for true navigation. The sensory basis underlying this feat is elusive, but presumably involves magnetic particles. A common experimental manipulation procedure consists of pre-treating animals with a magnetic pulse. This aims at re-magnetising particles to alter the internal representation of the external field prior to a navigation task. While pulsing provoked deflected bearings in laboratory experiments, analogous studies with free-flying songbirds yielded inconsistent results. Here, we pulsed European robins (*Erithacus rubecula*), being medium-distance migrants, at an offshore stopover site during spring migration and monitored their free-flight behaviour with a regional-scale tracking system. We found no pulse effect on departure probability, nocturnal departure timing, or departure direction, in agreement with results on a long-distance migrant released at the same site in autumn. This necessitates a reassessment of the importance of geomagnetic maps for migratory decisions for free-flying birds.

**Summary statement:** Magnetic pulse pre-treatment disturbs geomagnetic map usage of birds in lab environments. However, our free-flying birds show no effect, suggesting geomagnetic map information is less important in the natural environment.

## Introduction

Migratory songbirds navigate to familiar breeding and wintering sites with fascinating accuracy and precision, even over distances of several thousands of kilometres (e.g. Salewski et al. 2000; Holland et al. 2013). One cue that plays a major role in this navigational task is the Earth’s magnetic field. Through systematic changes in inclination, declination and total intensity, the Earth’s magnetic field forms a predictable grid around the globe (Skiles 1985) that could be used as a geomagnetic map. Thus, many studies on migrating birds have hypothesised that experienced birds use such a map to perform true navigation, defined as the ability to return to a familiar migratory destination from an unfamiliar location (reviewed in Holland 2014; Heyers et al. 2017). How birds detect geomagnetic information is still unknown, but is widely believed to be based on small magnetic particles innervated by sensory nerve endings of the trigeminal nerve (Kishkinev et al. 2013; Beason and Semm 1996). While we still lack direct anatomical evidence for the postulated magnetic particles (Curdt et al. 2022), their existence is supported by a number of behavioural studies that pre-treated birds with a strong magnetic pulse (e.g. Wiltschko and Wiltschko 1995; Holland and Helm 2013). The pulse is aimed at re-magnetising (Wiltschko et al. 1994) or re-arranging magnetic particles in the receptor (Davila et al. 2005), producing an altered sensory output to the unchanged ambient magnetic field. Thus, the magnetic pulse leads to an altered internal representation of the geomagnetic field for the bird. Under the magnetic map hypothesis, the pulsed animal would interpret such an altered magnetic field percept as a different location than present or as corrupted signal. Indeed, pulsed songbirds mostly showed deflected orientations, not only when tested in artificial orientation cages (Emlen-funnels; e.g. Wiltschko and Wiltschko 1995; Beason, Dussourd, and Deutschlander 1995; Wiltschko et al. 1998) but also when tracked in free-flight in the wild (Holland et al. 2009; Holland 2010; Holland and Helm 2013). Thereby, the magnetic pulse does not affect the magnetic compass of the birds, because the sensor is most likely based on a radical-pair-based mechanism containing no magnetic material (reviewed in Hore and Mouritsen 2016). Therefore, the results of the pulse studies are consistent with a magnetic-particle-based sensor in a geomagnetic map navigation context.

Using a regional high-throughput tracking system with unprecedented spatiotemporal resolution, we recently performed a pulse study comprising of 140 individuals of a long-distance migrant songbird, the northern wheatear (*Oenanthe oenanthe*, hereafter wheatear), on the offshore island of Helgoland in the German Bight during their autumn stopover (Karwinkel et al. 2022). To our surprise, pulsed birds and control birds turned out to be statistically indistinguishable in all migration-related behavioural decisions we had monitored, i.e., departure directions, departure probability, nocturnal departure timing, and consistency in flight direction over 50–100 km (Karwinkel et al. 2022). Taken all previous results together, the most parsimonious explanation for the absence of a pulse effect is the dispensability of magnetic map cues for the wheatear, at least at this offshore location. After all, the stopover site was still several thousands of kilometres away from the migratory destination in Africa, so magnetic map factors may not be essential for navigation at this particular part of the route. This rationale is supported by a recent ring-recovery study presenting correlative evidence that migratory birds use magnetic inclination angles as a cue when closing in on their target region (Wynn et al. 2022).

We consequently expanded our study to further explore the role of distance to final destination on magnetic map navigation and repeated our pulsing study with the same methodology on a short to medium distance migrant, the European robin (*Erithacus rubecula*, hereafter robin). During spring stopover on Helgoland, robins are much closer to their migratory destination [some 50 to 1250 km], in this case to their breeding sites (Dierschke et al. 2011), compared to wheatears in our former study, whose destination was their African wintering grounds (Karwinkel et al. 2022). Importantly, in an independently conducted study, free-flying robins have been reported to be affected by magnetic pulse pre-treatment on spring migration (Holland 2010). However, evidence to date was based on a sample size of only six treated and 13 control robins and a relatively short tracking range [maximum ca. 5.5 km] at a mainland location. Moreover, control and experimental birds were not corrected for physiological and environmental conditions (Holland 2010), meaning that important migratory factors, like energy stores of the birds, timing within the year and weather conditions differed between groups.

Consequently, here we investigate the response to the magnetic pulse of robins during spring migration on the following four migratory traits: departure probability, departure timing, departure direction and consistency in flight direction for the first 50–100 km after departure. We expect that pulsing has an effect on how birds extract and interpret geomagnetic map information, which in turn affects their decision of whether or not to depart, and if so, when and in what direction.

## Materials and methods

### Ethics

All work was conducted under the permission by the “Ministry of Energy, Agriculture, the Environment, Nature and Digitalisation” of the federal state Schleswig-Holstein, Germany under permission number V 244 – 16840/2019(41-4/19).

### Study site

The experiment was performed on Helgoland (54°11′ N, 07°53′ E), which is a small (about 1 km^2^) island in the North Sea (Fig. 1). The closest shorelines are on Wangerooge Island in the south (44 km) and at St. Peter-Ording (48 km; near Tümlauer Koog) in the east-north-east (Fig. 1B). The shoreline of the German Bight encompasses Helgoland in a section from about 17° (Sylt Island; 72 km) to 232° (Borkum Island; ∼ 104 km; Fig. 1B). Otherwise, Helgoland is surrounded by open sea, with the Norwegian coast about 425 km to the north and the British coast about 500 km to the west (Fig. 1A).

**Figure 1.**
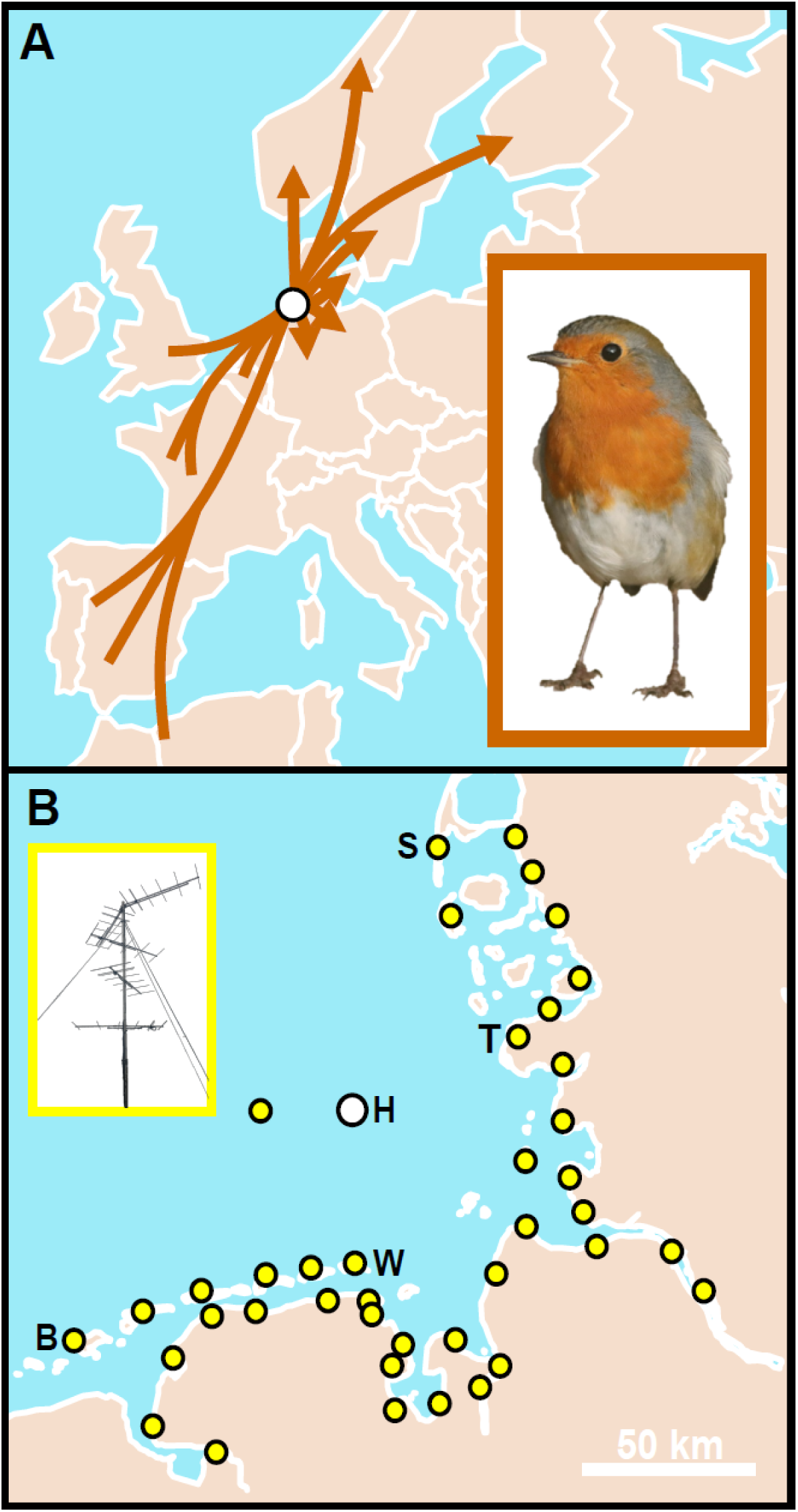
Study area in the context of European robin (*Erithacus rubecula*) migration and radio-tracking array. **(A)** Estimated migration routes of European robins passing Helgoland (H) on spring migration, based on ring recoveries. **(B)** Radio tracking array in the German Bight, with every dot representing a radio receiving station (example shown in the inset). H-Helgoland; B-Borkum; W-Wangerooge; T-Tümlauer Koog; S-Sylt.

### Study species

During spring migration, robins regularly stop over on the island, with about 200 individuals per day during peak passage from the end of March to the end of April (Dierschke et al. 2011). Robins are solitary night migrants, so their migratory behaviour is independent of conspecifics (Schmaljohann and Klinner 2020; Packmor et al. 2020; Dorka 1966). According to ring recoveries, robins passing Helgoland in spring migrate to northern Germany, southern Scandinavia and the Baltic region (Fig. 1A; Bairlein et al. 2014; Dierschke et al. 2011). The proposed breeding area is therefore about 50 to 1250 km away from our study side. Radio-tracking data of migrating robins from Helgoland show a mean departure direction of 90° (Klinner 2020). Robins are irregular breeders with maximum 1–2 pairs on Helgoland (Dierschke et al. 2011) and no breeding record in summer 2021 (J. Dierschke pers. comm.). Consequently, all robins included in our experiments were migrants. We caught robins with Helgoland funnel traps and mist nets between 24 March and 22 April 2021 throughout the day. Birds were aged, according to Svensson (1992) and Jenni and Winkler (2020). Only second-calendar year birds entered the experiment to exclude any potential age effects. Maximum wing chord of each individual was measured to the nearest 0.5 mm (Svensson 1992).

### Bird housing

After catching, birds were immediately transferred to individual cages (40 cm x 30 cm base, 40 cm high) within the island station of the Institute of Avian Research “Vogelwarte Helgoland”, where they received artificial light in the natural day-night rhythm. Birds were provided with *ad libitum* access to water and food (mealworms, *Tenebrio molitor*, supplemented with fat-food [Fett-Alleinfutter Typ II grün, Claus GmbH, Germany]) to allow for accumulation of fat, i.e. refuelling on energy stores. By that, we ensured that all birds carried sufficient and comparable energy stores important for high motivation to resume migration, as many studies have shown a significant positive correlation between energy stores and departure probability (Schmaljohann and Eikenaar 2017; Klinner et al. 2020; Eikenaar, Karwinkel, and Hessler 2021).

### Experimental treatment with a magnetic pulse

As night-migratory songbirds most likely decide to resume migration several hours before sunset (Eikenaar et al. 2020), we conducted the experiment six hours before sunset to provide pulsed birds with sufficient time to include the altered geomagnetic map percept into their migratory decision-making process. Pulsed birds that did not directly depart in the first night after treatment can be assumed to still have perceived an altered geomagnetic field over the following days, because former studies demonstrated effects for about ten days after magnetic pulsing (Wiltschko et al. 2007; Holland and Helm 2013). We conducted the experiments only on days with favourable migration conditions (wind <8 m s^-1^, no rain) to minimise weather-dependent effects on migratory decisions (Erni et al. 2002; Packmor et al. 2020). On these days, at about seven hours before sunset, we divided the housed robins into two equally-sized groups. To calculate the birds’ energy stores (Kelsey, Schmaljohann, and Bairlein 2020), we weighed them to the nearest 0.1 g and estimated their muscle score (Bairlein 1994). We found no difference in energy stores between the treatment and control group (Welch Two Sample t-test: t78=-0.852, p=0.315), which implies comparable physiological states between the groups.

Birds in the treatment group were exposed to a magnetic pulse of 0.1 T peak strength using the pulse geometry and protocol detailed in Karwinkel et al. (2022). In brief, the pulse was delivered with a small coil placed on the side of the beak and focused onto the middle part of the beak (Fig. 2), where the putative magnetic-particle-based sensor is thought to be located (Heyers et al. 2017). In the moment the pulse was fired, the birds were hand-held and their heads were fixed in a short piece of tubing (Fig. 2A) to ensure that the application geometry of the pulse was the same for each bird (Fig. 2). The head of the bird pointed southwards and the magnetic field lines of the pulse were aligned perpendicularly to the bird’s head, with the magnetic North pole pointing towards the bird (Fig. 2B). This perpendicular pulse field geometry was found to cause a significantly larger deflection of homing pigeons on the day of treatment compared to the axial pulse geometry (Beason, Wiltschko, and Wiltschko 1997).

**Figure 2.**
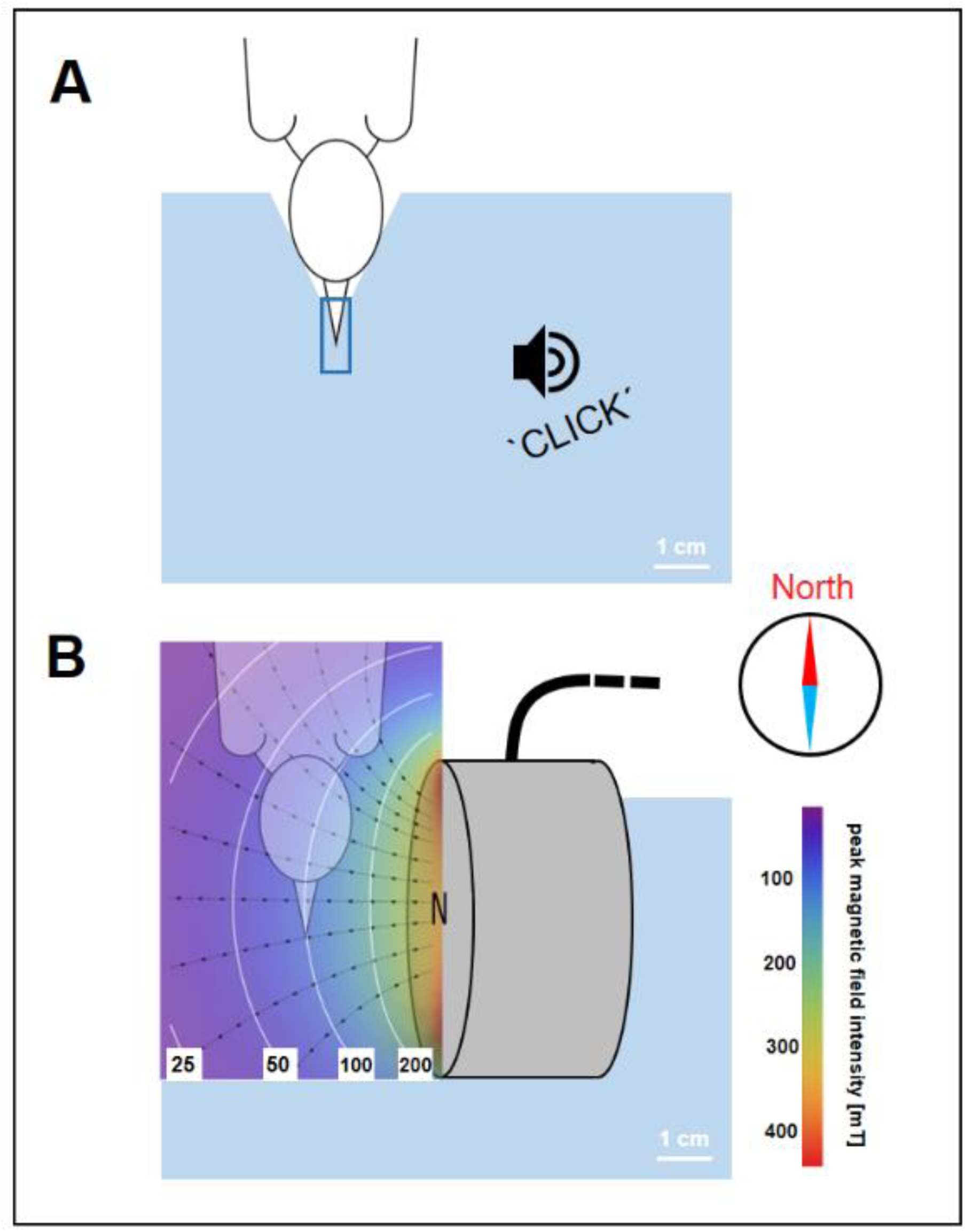
Control and experimental treatment with a magnetic pulse. **(A)** While hand-holding the bird, its head was gently fixed against an indentation on a styrofoam block, with its beak inserted in a small plastic tube (blue rectangle). Control birds experienced just a short ‘click’ sound. **(B)** Birds in the experimental group were similarly hand-held, but experienced a magnetic pulse from a coil (grey cylinder). White lines on the heat map show isolines of peak magnetic field intensity. Black lines are selected magnetic field lines. Figure adopted from Karwinkel et al. (2022).

The magnetic pulse was produced by a coil with a 5 cm diameter and 15 × 15 windings of 1 mm copper wire connected to a small generator (“Beck-Pulser”, magnetic pulse generator, Ing. Büro L. Albrecht, Heist, Germany). The magnetic field reached its maximum after 1.5 ms and decayed within 8 ms (Fig. S1). We ensured functionality of the pulse application at every experimental day by controlling peak pulse intensity with a magnetometer (Gaußmeter HGM09s, MAGSYS Magnet Systeme GmbH, Dortmund, Germany). The control group experienced the same handling procedure, but the magnetic pulse was replaced by a short ‘click’ sound produced by tapping a finger on the structure, resembling the sound of the magnetic pulser at firing (Fig. 2A). Afterward either treatment, we attached a radio-transmitter to the bird and released it before processing the next bird. We performed this experimental procedure on 31^st^ of March with 12 birds, on 20^th^ of April with 44 birds and on 25^th^ of April 2021 with 24 birds (n_total_=80). Each time, half of the birds entered the control group and the other half the treatment group.

### Radio-tracking free-flight behaviour

Birds were fitted with radio-transmitters (NTQB-2, Lotek Wireless Inc., Canada; burst intervals between 2.3 and 5.3 seconds) via individually adjusted leg-loop harnesses (Naef-Daenzer 2007). The total weight of the radio-transmitters including harness was about 0.31 g, which did not exceed 2.0% of the bird’s body mass (median: 1.8 %) and is therefore well below the 3–5% recommended threshold (Kenward 2001). To track the behaviour of the birds in free flight, we used an automated radio-receiving system on Helgoland (Müller et al. 2018; Karwinkel et al. 2022). This system consists of four SensorGnome receivers (www.sensorgnome.org) at three sites on Helgoland, connected to 16 radially aligned antennas (6-element Yagi antennas, Vårgårda Radio AB, Sweden), ensuring a radial resolution of 22.5° (Fig. S2A). Further, the German Bight is equipped with 40 comparable automated radio-receiving stations (Fig. 2B, Brust et al. 2019; Karwinkel et al. 2022), which allowed us to track the birds passing the shoreline of the German Bight after departure from Helgoland. All stations are part of the Motus Wildlife Tracking System; see http://www.motus.org and Taylor et al. (2017). From the radio-tracking data, as received from Motus, we determined (I) whether the bird stayed or departed in the first night after the treatment (departure probability), (II) the departure timing within the night, (III) departure direction from Helgoland and (IV) the consistency of this direction towards the shoreline (Fig. S2B), using an algorithm written by the authors, in a replicable and blinded approach to avoid any observer bias. For further details about the radio tracking data analysis see the supplemental material in Müller, Rüppel, and Schmaljohann (2018) and Packmor et al. (2020).

### Weather data

We considered weather parameters at two time points: First, weather parameters were assigned to the night after release at 200 minutes after sunset, which was the median departure timing in a former study with free-flying robins on Helgoland (Packmor et al. 2020), and second, for the individual time of actual departure.

Precipitation [mm] and cloud cover [eighths] were provided by a local weather station on the island operated by the German Weather Service (DWD). Airspeed flow assistance (hereafter wind assistance) [m s^-1^] for a flight direction of 41° (Klinner 2020), was calculated using NCEP reanalysis data (NOAA, Boulder, CO, USA, http://www.cdc.noaa.gov/cdc/data.ncep.reanalysis.derived.html; Kemp, Shamoun-Baranes, et al. 2012; Kemp, van Loon, et al. 2012). Wind assistance has the advantage of including multiple relevant wind parameters (e.g., side winds, tailwind component) as well as the birds’ own airspeed in a single unit.

### Statistics

All statistical analyses were performed using R 4.0.3 statistical software (R Core Team 2019). To assess whether our treatment affected the birds’ departure probability in the first night [departing or not departing, binomial], we ran a generalised linear model (GLM) including the following predictor variables: treatment condition [experiment/control; categorical], wind assistance [continuous], cloud cover [categorical] and energy stores [continuous]. Here, weather parameters were considered from 200 minutes after sunset for the night after release (see section above). We z-transformed all numeric variables. Since model assumptions were violated, as detected by goodness of fit plots (Korner-Nievergelt et al. 2015), and since we did not find a transformation to solve this problem, we had to discard the model and ultimately used a chi-square test to assess the potential difference in the departure probability between the control and experimental group.

Because nights were getting shorter during the course of the experiment (night duration of first experiment: 664 min; night duration of last experiment: 558 min), to determine an individual’s departure timing, we calculated the proportion of night at departure for each individual separately. To do so, an individual’s departure time [minutes after sunset] was divided by the night length in the night of departure [minutes]. To explain variation in the departure timing [log-transformed], we applied a linear model (LM) including the following predictor variables: treatment condition [experiment/control; categorical], wind assistance [continuous], cloud cover [categorical], day of year of release [Julian Day, see above], and energy stores [continuous] and all two-way interactions with the treatment condition (all numerical variable z-transformed). Weather parameters and day of year were assigned for the individual departure timing. Predictor variables were found to be linearly independent (VIF < 2; Babak 2013). Since none of the two-way interactions came out as significant, they were removed from the final model. The residual analyses did not indicate violation of model assumptions (see R-Code in the supplemental for details).

To compare the circular variables (departure direction, consistency in flight direction) between the groups, we applied circular statistics using the R packages ‘CircStats’ (Lund and Agostinelli 2012) and ‘circular’ (Lund and Agostinelli 2013). Due to the arrangement of the radio-tracking antennas on Helgoland (Fig. S2A), our directional data was grouped. Since the Mardia-Watson-Wheeler test randomly breaks ties (in our case the groups), we repeated the test 10,000 times to exclude any bias due to random tie-breaking, and then provided the median of the test parameters (see R-Code in the supplemental for details).

## Results and Discussion

### Effects on the four migratory traits

Departure probability was 83% in the control group (33 out of 40 robins departed in the first night after release) and 90% in the magnetic-pulse treated group (36 out of 40 departed in the first night, Fig. 3A, Table S1). Our results thus fail to demonstrate an effect of the magnetic pulse, supposed to alter geomagnetic map information, on the motivation to migrate as departure probability between the groups did not differ (Pearson’s χ^2^-test: χ^2^_1_=0.42, p=0.52). A former study demonstrated that artificially altered geomagnetic field information on caged birds indeed affected nocturnal migratory restlessness (Bulte et al. 2017), which is a direct proxy for the departure probability in the wild (Eikenaar et al. 2014). However, while Bulte et al. (2017) altered the geomagnetic field consistent with a meaningful spatiotemporal occurrence of the study species, we corrupted the sensor of the robins with a magnetic pulse in an unknown way. This makes it impossible to predict, whether the field percept corresponds to a meaningful location to our birds or not. Nevertheless, the study of Bulte et al. (2017) shows that changes in geomagnetic cues affect the motivation to depart in caged birds, but our results and the former ones of Karwinkel et al. (2022) did not verify this for free-flying songbirds.

**Figure 3.**
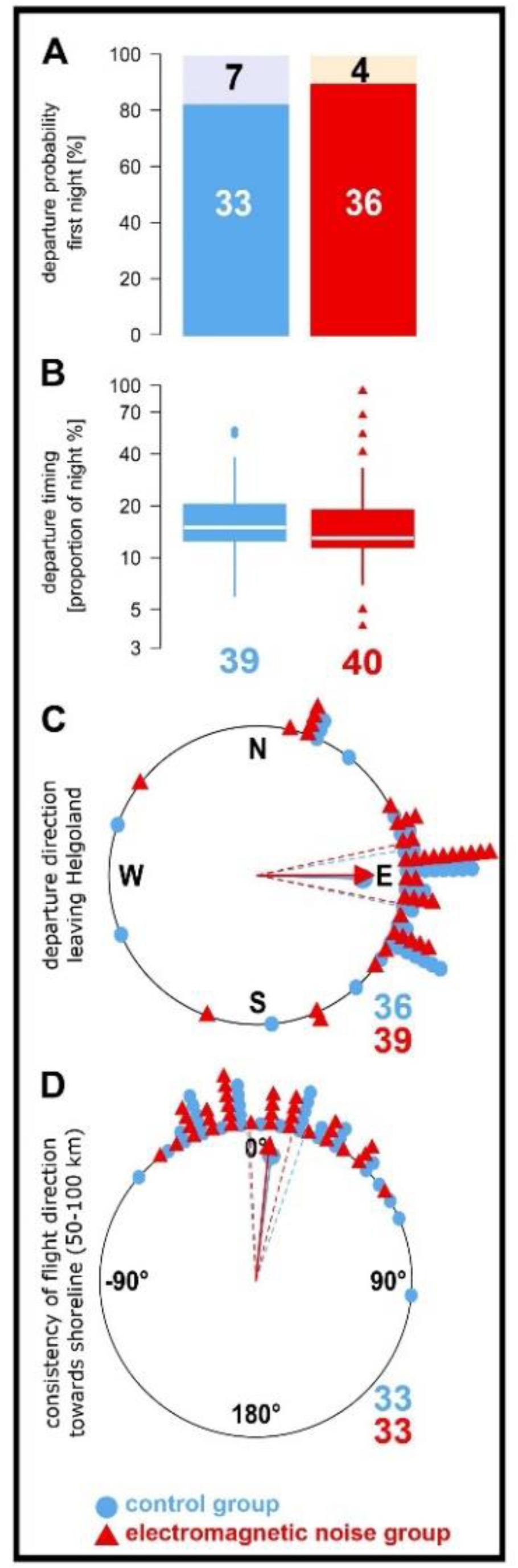
Effect of a magnetic pulse on four migratory traits in free-flying European robins (*Erithacus rubecula*) during spring migration. Numbers indicate sample sizes per group. Sample sizes decreased in sequence as not every trait could be characterised for every bird, for details see methods. **(A)** Departure probability as proportion of birds departing on the first night after the release from Helgoland (white numbers, lower bar solid colour) or staying for the first night (black numbers, upper bar transparent colour). **(B)** Nocturnal departure timing as proportion of the night at departure for all birds. Control: 1st quartile: 13%, median: 15%, 3rd quartile: 21%, range: 6–55%. Treatment: 1st quartile: 12%, median: 13%, 3rd quartile: 19%, range: 4–93%). **(C)** Initial departure direction from Helgoland. Control: Rayleigh-test: r=0.77, p < 0.001, mean: 92°. Treatment: Rayleigh-test: r=0.78, p < 0.001, mean: 90°. **(D)** The consistency of flight direction after departure from Helgoland until passage at the shoreline of the German Bight (50–100 km), given as the directional deviation between departure direction from Helgoland and passage location site on the shoreline (Fig. 1B, see methods for more details). Control: Rayleigh-test: r=0.856, p<0.001, mean: 7°. Treatment: Rayleigh-test: r=0.919, p<0.001, mean: 6°. Data points in the circular plots are shifted slightly off-centre by <5° to better distinguish the data of the corresponding groups.

Regarding nocturnal departure timing, robins in the control group had a median departure after 15% of the night length had elapsed, and robins of the magnetic-pulse treated group after 13% had elapsed (Fig. 3B). We found no effects of the treatment condition (p=0.64; Fig. 3B) and day of year (p=0.64) on nocturnal departure timing (Table S2). Increasing energy stores (p=0.03) and wind assistance (p < 0.001), but decreasing cloud cover (p=0.002), advanced nocturnal departure timing (Table S2), which is consistent with the natural departure behaviour of free-flying birds, i.e., without caging and feeding (e.g. Packmor et al. 2020). This reflects that we can confidently assume that our experimental procedure does not alter the natural migration behaviour of robins in this study.

Departure directions from Helgoland of both groups were significantly oriented to the east from Helgoland (Fig. 3C) and did not differ from each other (repeated Mardia-Watson-Wheeler test: W_2_=0.33, p=0.85). The robins consistently maintained their flight direction from Helgoland of about 50–100km towards the shoreline (Fig. 3D) and their consistency did not differ between the groups (Watson-Williams test: F_1,64_=0.04, p=0.842). Consequently, departure directions from Helgoland are representative for the sustained migratory flight direction during the night. Therefore, we are convinced that there was no biologically significant effect of the magnetic pulse on departure direction. This is in stark contrast to the findings of (Holland 2010), who found a 90° anticlockwise shift in departure direction after a magnetic pulse applied to the same species, also during spring, albeit with a fifth of our sample size.

### Is geomagnetic map information dispensable for migratory birds?

Former magnetic pulse studies (e.g. Wiltschko and Wiltschko 1995; Holland and Helm 2013) suggested that birds use geomagnetic map information obtained via a magnetic-particle-based sensor. This is in contrast to our findings where, despite large statistical power to resolve effects if present, pulse pre-treatment did not affect migratory behaviour and decisions, neither in robins as reported here, nor in our parallel study on wheatears (Karwinkel et al. 2022). We can safely assume that our pulse was strong enough to affect the putative magnetite-based magnetoreceptor and to provoke altered magnetic field percepts persisting over the observation time. It is also evident that the departure directions of our tracked birds are seasonally appropriate. However, a key requirement for the magnetic pulse to have an effect on migratory decisions is that birds rely on magnetic map information for their migratory decisions. In former studies, this was assumed, although it was mostly unknown before the experiments were performed. This prompts rethinking of the necessity of geomagnetic map cues for the migratory decisions considered in our study.

The assumed accuracy of a geomagnetic map is supposed to be in the order of 10–30 km (Mouritsen 2018). As all robins left Helgoland and migrated at least beyond the shoreline of the German Bight, their breeding areas are most likely far more than 50–100 km away from the island. Thus, the spatial resolution of the geomagnetic map information is deemed sufficient to guide the robins from Helgoland towards their breeding areas (Mouritsen 2018). However, during which parts of their migratory journey birds rely on geomagnetic map information for navigation and where they may dispense with it, is currently unclear. Former studies that found an effect might have been conducted at locations critical for geomagnetic map navigation, for example, if the birds were in the homing range of the migratory goal (Holland 2010; Holland and Helm 2013). Birds stopping over on Helgoland may be in a different phase of migration, e.g., long-distance phase. Thus, the importance of geomagnetic map cues for deciding in which direction to continue migration might differ between the studies in general and might be rather low in our case in particular.

It appears most likely that birds (both pulsed and untreated), independent of their migration context, rely primarily on non-magnetic map cues or no map cues at all in places like Helgoland. Of all cues thought to be important for navigation in migrants, landmarks and similar visual patterns are considered to be particularly important for songbirds (Mouritsen 2018). Since most migratory songbirds show weak faithfulness to their stopover sites (Catry et al. 2004), which is especially true for Helgoland (Dierschke 2002), our birds were unlikely to have former experience about the specific location of Helgoland. However, general landmarks in the German Bight, such as the shoreline or artificially illuminated areas, might guide them towards a given location. On the other hand, comparable landmarks, e.g., mountain ridges, rivers, Lake Constance, streets or cities, were also present and even far closer in the studies reporting pulse effects (Holland 2010; Holland and Helm 2013). Therefore, it is not obvious why such landmarks would be more important at our test site than at others. It remains unclear why free-flying birds in Holland (2010) and Holland and Helm (2013) responded to the magnetic pulse with a deflected departure direction, but not in our present study or in Karwinkel et al. (2022).

The contradictory findings on geomagnetic map information in birds discussed above warrant rethinking the perception and use of geomagnetic maps for migration decisions in a sensory and ecological context. To better understand or even resolve this contradiction, a magnetic pulse study is needed where we know with certainty that birds rely on and indeed use geomagnetic map cues, as shown in Chernetsov, Kishkinev, and Mouritsen (2008) or Kishkinev et al. (2015). Such a study would ultimately link the geomagnetic map use and to a sensory system based on magnetic particles.

### Do birds need a magnetic-particle-based mechanism for geomagnetic map navigation?

If an experiment like the one suggested above does not find an effect of a magnetic pulse, we carefully question that a magnetic-particle-based mechanism is the sensory mechanism required for geomagnetic map navigation. Of the three geographic properties of the Earth’s magnetic field (inclination, declination, and total intensity), the radical-pair-based mechanism of magnetoreception might be able to sense at least inclination and declination (declination only in combination with a celestial compass). Whereas there is direct evidence for the use of declination (Chernetsov et al. 2017), and partly also for inclination (Wiltschko and Wiltschko 1992; Wynn et al. 2022) as geomagnetic cues, convincing evidence for the biological importance of magnetic intensity for navigation is currently lacking. This at least raises the cautious question of whether birds need a magnetic-particle-based mechanism for geomagnetic navigation and therefore whether such a sensor exists.

## Acknowledgements

We thank the team at the island station of the Institute of Avian Research on Helgoland for their support during fieldwork, especially Sven Hessler, Jochen Dierschke and Klaus Müller. We thank Mario de Neidels, Thomas Mertens, Heinz-Hinrich Blikslager and the entire team behind the Motus wildlife tracking system for their invaluable technical support. We also thank Dario Allenstein for the independent radio tracking data analysis, Zephyr Züst for language editing and Miriam Liedvogel for helpful comments.

## Authors’ contributions

H.S., F.B., and M.W. conceptualisation and methodology; T.K., and E.J.: field work; T.K., E.J., and H.S.: data analyses and statistics; H.S., O.H., and V.B.: Motus radio-tracking system; T.K. and H.S.: writing-original draft; all: revising and editing of the manuscript; H.S.: supervision. All authors gave their final approval for publication and agree to be held accountable for the work performed therein.

## Data availability

The datasets and codes supporting this article have been uploaded as part of the supplementary material.

